# A contamination focused approach for optimizing the single-cell RNA-seq experiment

**DOI:** 10.1101/2022.10.25.513758

**Authors:** Deronisha Arceneaux, Zhengyi Chen, Alan J. Simmons, Cody N. Heiser, Austin N. Southard-Smith, Michael J. Brenan, Yilin Yang, Bob Chen, Yanwen Xu, Eunyoung Choi, Joshua D. Campbell, Qi Liu, Ken S. Lau

**Author notes:** These authors contributed equally (Deronisha Arceneaux, Zhengyi Chen). Correspondence (KSL), Ken S. Lau, Associate Professor of Cell and Developmental Biology, Vanderbilt University School of Medicine, 2215 Garland Ave, 10405A MRB IV, Nashville, TN 37232-0441.

## Abstract

Achieving high data quality in single-cell RNA-seq (scRNA-seq) experiments has always been a significant challenge stemming from minute signal that can be detected in individual cells. Droplet-based scRNA-seq additionally suffers from ambient contamination, comprising nucleic acid materials released by dead cells into the loading buffer and co-encapsulated with real cells, which further washes out real biological signals. Here, we developed quantitative, ambient contamination-based metrics and an associated software package that can both evaluate current datasets and guide new experimental optimizations. We performed a series of experimental optimizations using the inDrops platform to address the mechanical and microfluidic cell encapsulation aspect of an scRNA-seq experiment, with a focus on minimizing ambient contamination. We report improvements that can be achieved via cell fixation, microfluidic loading, microfluidic dilution, and nuclei versus cell preparation; many of these parameters are inaccessible on commercial platforms. We provide insights into previously obscured factors that can affect scRNA-seq data quality and suggest mitigation strategies that can guide future experiments.

## Introduction

Single-cell RNA sequencing (scRNA-seq) is a technique that allows for the investigation of genome-scale gene expression in thousands of individual cells, facilitating the deconvolution of tissue heterogeneity and population dynamics. The most popular scRNA-seq platforms involve microfluidic encapsulations of cells and barcoded capture oligonucleotides in oil emulsions, that ultimately enables sequencing reads to be assigned to each droplet or cell (Klein et al., 2015; Macosko et al., 2015; Zheng et al., 2017). Because of the low cellular loading required to avoid two or more cells captured in an individual droplet, most droplets are devoid of cells and ideally only contain loading buffer and RNA-capture beads. However, during the cell handling process, cell death, lysis, and leakage results in ambient RNA deposited into the loading buffer; this RNA is either co-captured with cells into droplets or into empty droplets themselves (Young and Behjati, 2020). Ambient RNA contamination lowers effective sequencing read depth, and more importantly, contributes to an insidious signal that masks true biological signals and confounds downstream biological interpretation.

To reduce cellular stress and enhance cell viability, a variety of tissue dissociation protocols has been developed, many of which were optimized for specific tissues and cell types. For instance, van der Wijst and colleagues developed a one-step collagenase dissociation protocol to combat batch effects introduced during the handling of cryopreserved gut mucosal biopsies (Uniken Venema et al., 2022). A listing of many of these protocols matched to tissue types was presented by Regev and colleagues (Slyper et al., 2020). Many of these protocols generate highly viable cells coming out of dissociation, as assessed by flow cytometry and live/dead dye visualization. However, they do not address the continuous stresses that cells are exposed to downstream of dissociation but prior to lysis in encapsulated droplets. Single cells in suspensions removed from their native tissue niches are often prone to cell death (Vachon, 2018). Single-nucleus RNA-seq (snRNA-seq) has been developed for difficult-to-dissociate tissues (e.g., neuronal tissues) (Lacar et al., 2016). Since nuclei are not cells, the prevalent thought in the field is that they are more resistant to physical stresses when isolated. However, substantial amounts of cytoplasmic RNA and ribosomes adhere to the surfaces of isolated nuclei (Slyper et al., 2020), and their contribution to downstream data analysis is undefined. Furthermore, the ‘gold standard’ for assessing protocol effectiveness has been the qualitative evaluation of genes and cell types recovered in the resulting data. Their effects on ambient contamination have not been directly nor quantitatively assessed.

In this study, we conduct controlled experiments to assess cell handling and microfluidics encapsulation variables that contribute to cell death and ambient RNA contamination. We developed quantitative contamination-based metrics to assess ambient RNA encapsulated into droplets as reflected in downstream sequencing data. We use the inDrops open scRNA-seq platform to manipulate microfluidics parameters that are inaccessible to commercial systems (Klein et al., 2015). We demonstrate our systematic evaluation of cell handling and microfluidics parameters that affect ambient contamination. We also provide quantitative and qualitative methods to deduce data quality upstream of data analysis that can potentially alert biologists to hidden quality control issues within data.

## Results

### Metrics that focus on ambient contamination can identify poor quality scRNA-seq datasets

We set out to first develop a set of quantitative metrics focused on ambient RNA levels, such that modifications made to downstream protocols can be adequately assessed. For illustrating situations with high and low ambient contamination, we used CellBender (Fleming et al., 2022) to simulate representative high quality (ambient gene count =100) and low quality (ambient gene count = 4000) datasets, while keeping other parameters relatively constant **(Fig. S1A-D)**. Since ambient contamination is present in real cells as well as empty droplets, we developed metrics that apply to unfiltered data. This way, ambient contamination assessment can be performed automatically without the subjectivity of data filtering.

Ambient RNA disrupts the ability to separate real cells from empty droplets, as can be illustrated by the standard UMI count versus log ranked barcodes curve **(Fig. S1E-F)**, which can also be represented as the cumulative distribution of counts versus ranked barcodes **(Fig. S1G-H)**. We scaled the total number of barcodes analyzed to the number of expected cells for each dataset to enable comparison between samples with different numbers of encapsulated cells and empty droplets. A sharp change in slope of the cumulative count curve, with a clear inflection point, was observed in the high-quality dataset because real cells contribute to notably larger increments of gene counts than empty droplets when background noise is low. The change in slope was less apparent in the low-quality dataset because ambient genes contribute to high increments of gene counts in empty droplets **(Fig. S1G-H)**.

The ability to separate true signals from background noise can be assessed in two ways: geometrically or statistically. Geometrically, a cumulative count curve resembling a rectangular hyperbola reflects this sharp change in slope, and hence higher quality, while the resemblance to a straight line reflects the opposite. For quantification, we defined secant lines connecting each point on the cumulative count curve to the diagonal line linking the origin to the last data point of the cumulative count curve **(Fig. 1A-B)**. The high-quality dataset, due to its resemblance to a rectangular hyperbola, has a larger maximal secant line distance as well as a larger standard deviation over all secant line distances, as compared to the low-quality dataset **(Fig. 1A-B)**. We also assessed the direct resemblance of the curve to a rectangle by calculating an area ratio between the area under the cumulative count curve and the minimal rectangle circumscribing the curve, which we termed AUC percentage over minimal rectangle, with high quality data occupying more of the rectangular area **(Fig. 1A-B)**. We then inverted these quantitative assessments to establish contamination metrics, such that they scale in proportion to the degree of contamination according to the geometry of the cumulative counts versus ranked barcodes curve **(Fig. 1A-B)**.

**Fig. 1.**
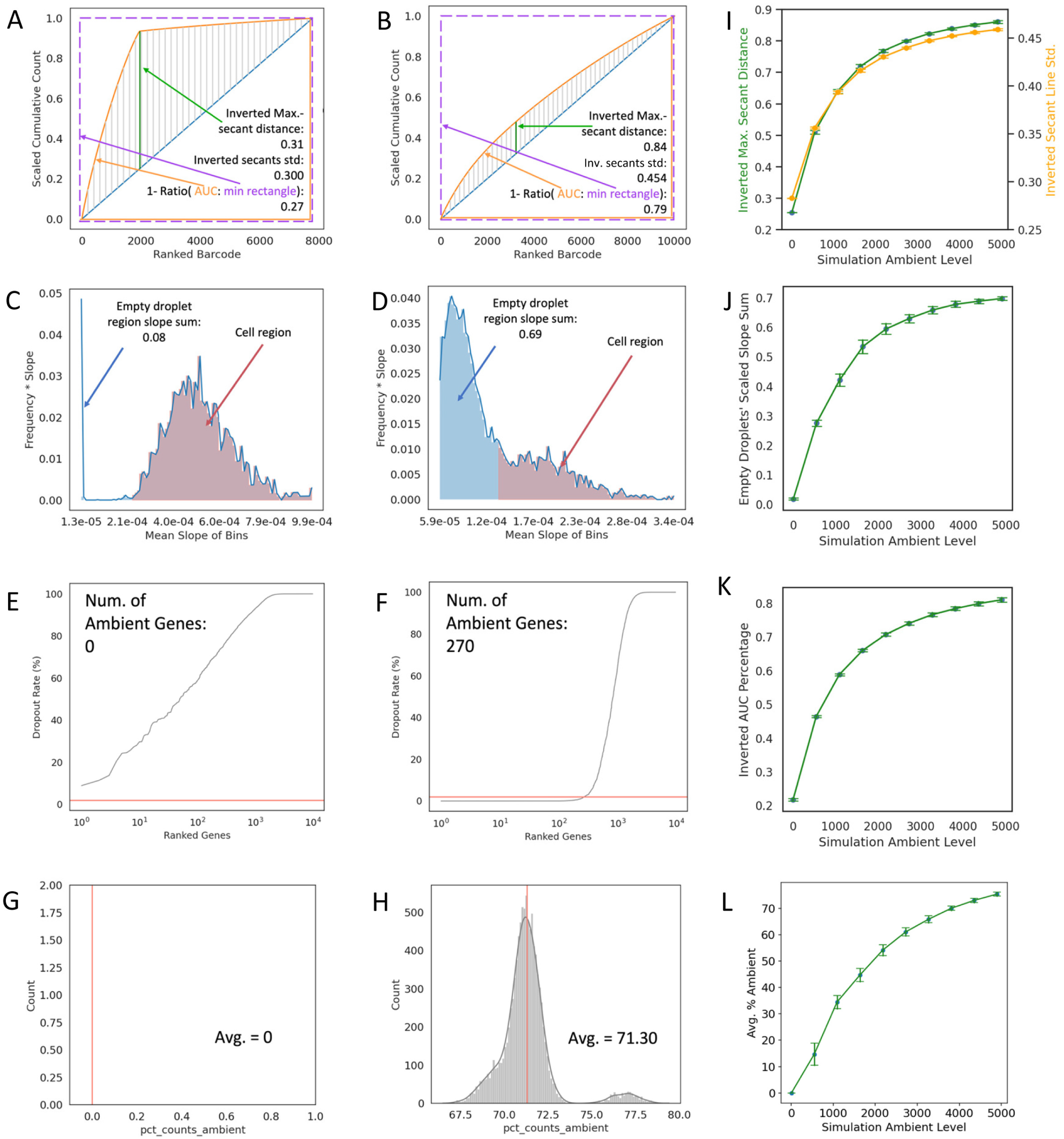
Ambient contamination metrics robustly reflect data quality on simulated datasets. **(A-B)** Scaled cumulative total transcript counts over ranked barcodes by total transcript counts for datasets simulated with **(A)** low ambient level and **(B)** high ambient level. Secant lines from the curve to the diagonal line are colored in grey with the line with maximal secant line colored in green, which were used to calculate inverted maximal secant distance and secant line standard deviation. The area under curve (colored in orange) and the minimal rectangle circumscribing (dashed purple line) were used to calculate the inverted AUC percentage. **(C-D)** Scaled frequency over slope for curves shown in **(A-B)**, respectively. The y-axis values were calculated by scaling the frequency to the mean slope per histogram bins on the x-axis. The region representing slopes that were below the threshold to be considered empty droplets were colored in blue and was quantified as empty droplets’ scaled slope sum. **(E-F)** Distribution of dropout rate of genes ranked by ascending dropout rate for datasets simulated with **(E)** low and **(F)** high ambient level. The pink line is drawn at 2% dropout rate, the cut-off below which a gene will be defined as ambient. **(G-H)** Distribution of percentage of ambient genes expressed per cell for dataset simulated with **(G)** low and **(H)** high ambient level. The mean percentage is quantified. **(I-L) (I)** Maximal secant distance (green) and secant line standard deviation (yellow), **(J)** AUC percentage, **(K)** cell’s scaled slope sum, and **(L)** percent counts ambient over different ambient levels for simulations. Line plots shown as mean± stdev of n=1000 replicates for each ambient level.

We also used statistical distributions to quantify ambient contamination, by first generating a distribution of slopes at each point of the cumulative count curve **(Fig. S1I-J)**. By further scaling each bin of the histogram with the mean of each bin’s x-axis slope span, the heights of bins capturing data points with high increments of gene counts, thus, from real cells, were scaled up and can be visualized **(Fig. 1C-D)**. The scaled distributions were normalized to one to enable cross dataset comparisons. We surmised that a contaminated dataset should have a slope distribution closer to unimodal, due to indistinguishable cells and empty droplets, while a high-quality dataset should have a multimodal slope distribution. Thus, a cut-off was determined to separate an “empty droplet” slope distribution from a “cell” slope distribution. This cut off was determined to be one standard deviation above the median of all slopes to approximate the “empty droplet” distribution, since most barcodes are empty droplets in a scRNA-seq experiment **(Fig. 1C-D)**. The sum of scaled slopes below this threshold, denoting data points that are potentially background ambient signals, is a quantitative metric that scales with dataset’s contamination level **(Fig. 1C-D)**. Aside from slope distributions, we further constructed distributions of ambient genes **(Fig. 1E-F)**. These are genes detected to be present in most barcodes (both cells and empty droplets) and have a dropout rate of less than 2% (Heiser et al., 2021). The number of ambient genes and the mean percentage of the ambient gene expressed per cell quantitatively differed between high- and low-quality datasets **(Fig. 1E-H)**. Thus, summary metrics from various statistical distributions can also be used to quantitatively assess ambient contamination.

To verify the robustness of these metrics to quantitatively assess ambient contamination, we evaluated simulated datasets at 10 ambient levels over n=1000 replicates. The contamination metrics - inverted maximal secant distance, inverted secant line standard deviation, inverted AUC percentage, sum of weighted slopes under threshold, average percentage of ambient genes, and the number of ambient genes all quantitatively increased in proportion to the ambient level set **(Fig. 1I-L; Fig. S1K)**. These results demonstrate that the quantitative metrics derived, which we termed contamination metrics, can robustly inform scRNA-seq data quality on a continuous scale based on ambient RNA contamination.

### Application of contamination metrics revealed ambient contamination in datasets that passed QC

We applied contamination metrics on datasets generated from the inDrops platform without any modification (standard inDrops)(Klein et al., 2015). K562 cells, as optimized in the original inDrops manuscript, demonstrated low contamination based on our metrics (Sample 1 - Inverted Max. Secant Distance: 0.28, Inverted Secant Line St. Dev.: 0.282, Inverted Percentage AUC: 0.20, Empty Droplet’s Scaled Slope Sum 0.17, Avg. Percent Counts Ambient: 5.5; Sample 2-Inverted Max. Secant Distance: 0.39, Inverted Secant Line St. Dev.: 0.315, Inverted Percentage AUC: 0.3, Empty Droplet’s Scaled Slope Sum: 0.15, Avg. Percent Counts Ambient: 0; Sample 3-Inverted Max. Secant Distance: 0.34, Inverted Secant Line St. Dev.: 0.301, Inverted Percentage AUC: 0.26, Empty Droplet’s Scaled Slope Sum: 0.11, Avg. Percent Counts Ambient: 0), as well as standard QC metrics such as mitochondrial count percentage per cell **(Fig. 2A-D; Fig. S2A-K; Table S1)**(Hong et al., 2022). Cultured cells maintain very high viability after minimal or no dissociation, leading to high data quality. In contrast, we also dissociated gastric corpus tissues in an unoptimized fashion and applied inDrops (see STAR Methods). The gastric corpus is the site of stomach acid production and houses various types of gastric cells, including acid-producing parietal cells (Engevik et al., 2020). Thus, dissociated single cells in this environment are exposed to extrinsic stress and damage. Standard data analysis revealed obvious QC failure in scRNA-seq data generated, as reflected by a high mitochondrial percentage, low number of genes detected, and general inability to detect known cell types **(Fig. S2L; Table S1)**. Poor data quality was also captured by our contamination metrics **(Fig. 2E-H; Table S1)**. While obvious QC failure is easy to detect, there are intermediate cases where low data quality can be masked within data that qualitatively passed QC, such as the case with the colonic epithelium. The colonic epithelium consists of an unilaminar layer of connected differentiated and undifferentiated epithelial cells. Differentiated cells do not self-renew and undergo anoikis when dissociated from their neighbors, increasing the propensity of dying cells in suspension (Vachon, 2018). More importantly, secretory cells, including goblet cells, are constantly under endoplasmic reticulum stress (due to heightened protein production) and are packed with tubulovesicular elements for protein secretion, leading to increased fragility (Kaser and Blumberg, 2009). Standard inDrops scRNA-seq of colonic epithelium (Southard-Smith et al., 2020) did not lead to QC failure, and two major lineages of secretory and absorptive cells can clearly be delineated from the data **(Fig. S2M; Table S1)**. However, closer examination of the data revealed that the high expressing, Goblet cell-specific gene *Muc2* was found in every cell, demonstrating a high degree of ambient contamination **(Fig. S2N)**. Standard quality metrics such as mitochondrial count percentage, total UMI count, and total genes were unable to distinguish datasets of low versus high quality arising from ambient RNA, but our contamination-based metrics could **(Fig. 2I-L; Table S1)**. These results demonstrate that our contamination metrics can reveal previously missed problems of ambient RNA in data generated using droplet-based scRNA-seq.

**Fig. 2.**
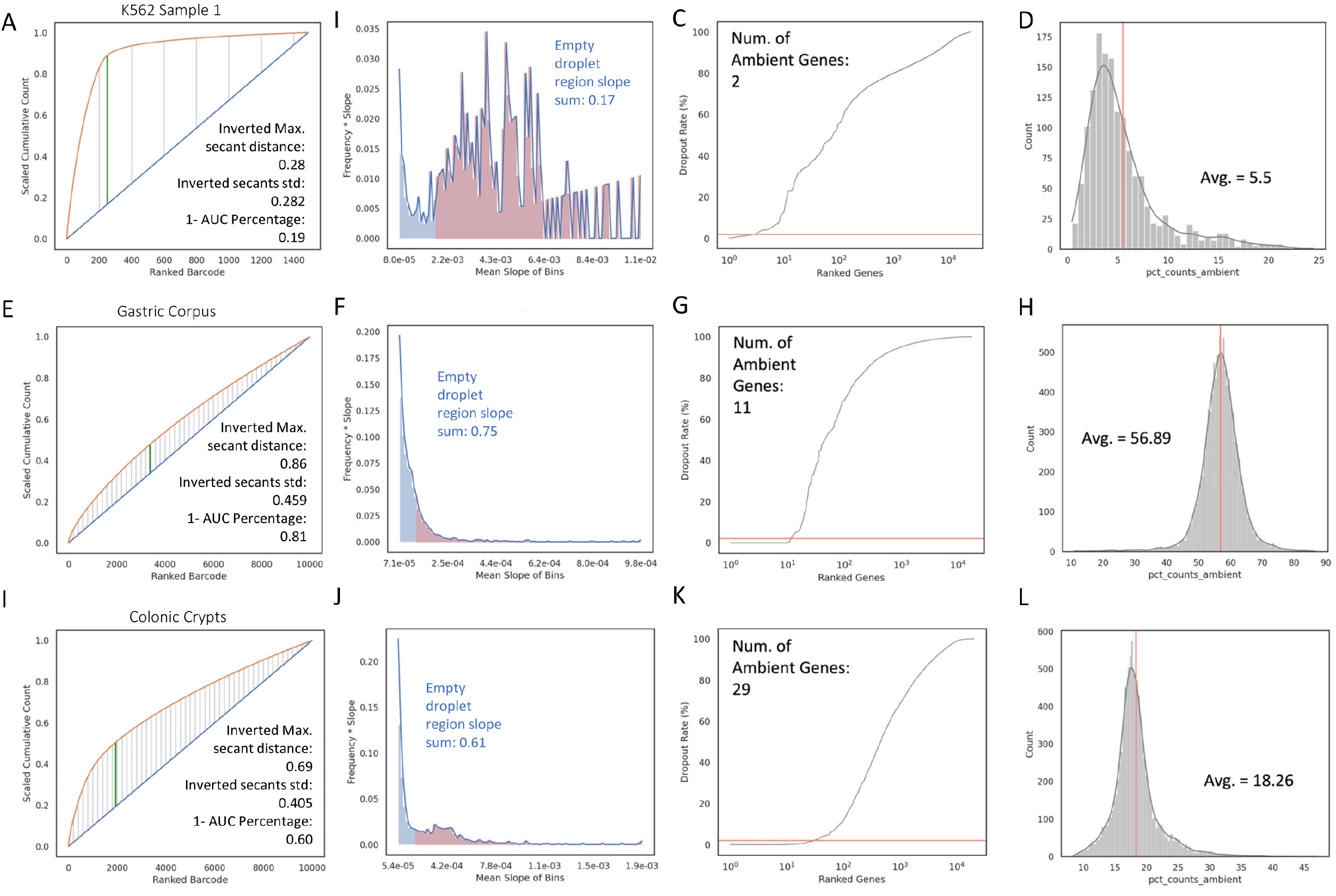
Contamination metrics on experimental datasets inform data quality on a continuous scale. Ambient contamination plots and metrics, formatted similarly to **Fig. 1** of experimental datasets of different quality: **(A-D)** K562 (Sample 1) cell line, **(E-H)** mouse gastric corpus, **(I-L)** and mouse colonic crypts.

### Pre-encapsulation variables affect scRNA-seq data quality and cell type diversity

Using colonic epithelium as our model system, we sought pre-encapsulation variables that may impact downstream ambient contamination. Slyper et al. revealed the impact of dissociation strategies on scRNA-seq profiling of various tissue types (Slyper et al., 2020). We first compared scRNA-seq data quality following standard total tissue dissociation protocols (https://dx.doi.org/10.17504/protocols.io.busfnwbn) compared with our standard crypt isolation strategy followed by cold protease dissociation as a control (cold protease and DNase cocktail; see STAR Methods). Protocols that started with total tissue mincing resulted in extremely high levels of ambient RNA, leading to near QC failure results **(Table 1; Fig. S3A-B)**. We surmised that the act of mechanical separation led to tissue stress and cell damage, leading to unacceptable cell death. This cell death can be clearly visualized in the cell hopper of the microfluidic encapsulation chip **(Fig. 3A)**. Thus, we proceeded to test different single-cell dissociation enzymes on chelated crypts to minimize tissue trauma. To our surprise, standard dissociation enzymes, collagenase, and DNase, and the Miltenyi MACs enzyme also led to high contamination, resulting in near QC failure **(Table 1; Fig. S3C-D)**. These dissociation enzymes require long incubation times at 37°C which may accelerate biological processes including cell death in single-cell suspensions, while cold protease has been shown to preserve viability by the opposite effect (Adam et al., 2017). Contamination metrics calculated on cold protease dissociation on crypts datasets were quantitatively lower than near QC failure datasets **(Fig. 3B-C; Figure S3E-F)**. The percent count ambient metric was more variable since the identities of ambient genes differed amongst techniques **(Fig. 3D)**. Standard QC metrics, such as number of genes and transcripts detected, also demonstrated higher data quality derived from cold protease dissociation on crypts compared to conditions that lead to near QC failure **(Fig. S3G-J)**. We also assessed the effect of dissociation enzymes on cell type representation, with the null hypothesis of negligible differences. We compared short term TrypLE dissociation on colonic crypts, which also resulted in decent data quality, to cold protease dissociation. Integrated UMAP and clustering analysis indicated that TrypLE resulted in recovering more immune cells, while cold protease recovered more tuft cells and more cells in general **(Fig. 3E)**. While standard dissociation strategies can result in acceptable data quality with hardy cell types such as cancer cells, our results demonstrate how various stressors at the tissue and the cell level can affect downstream ambient contamination and cell recovery.

**Fig. 3.**
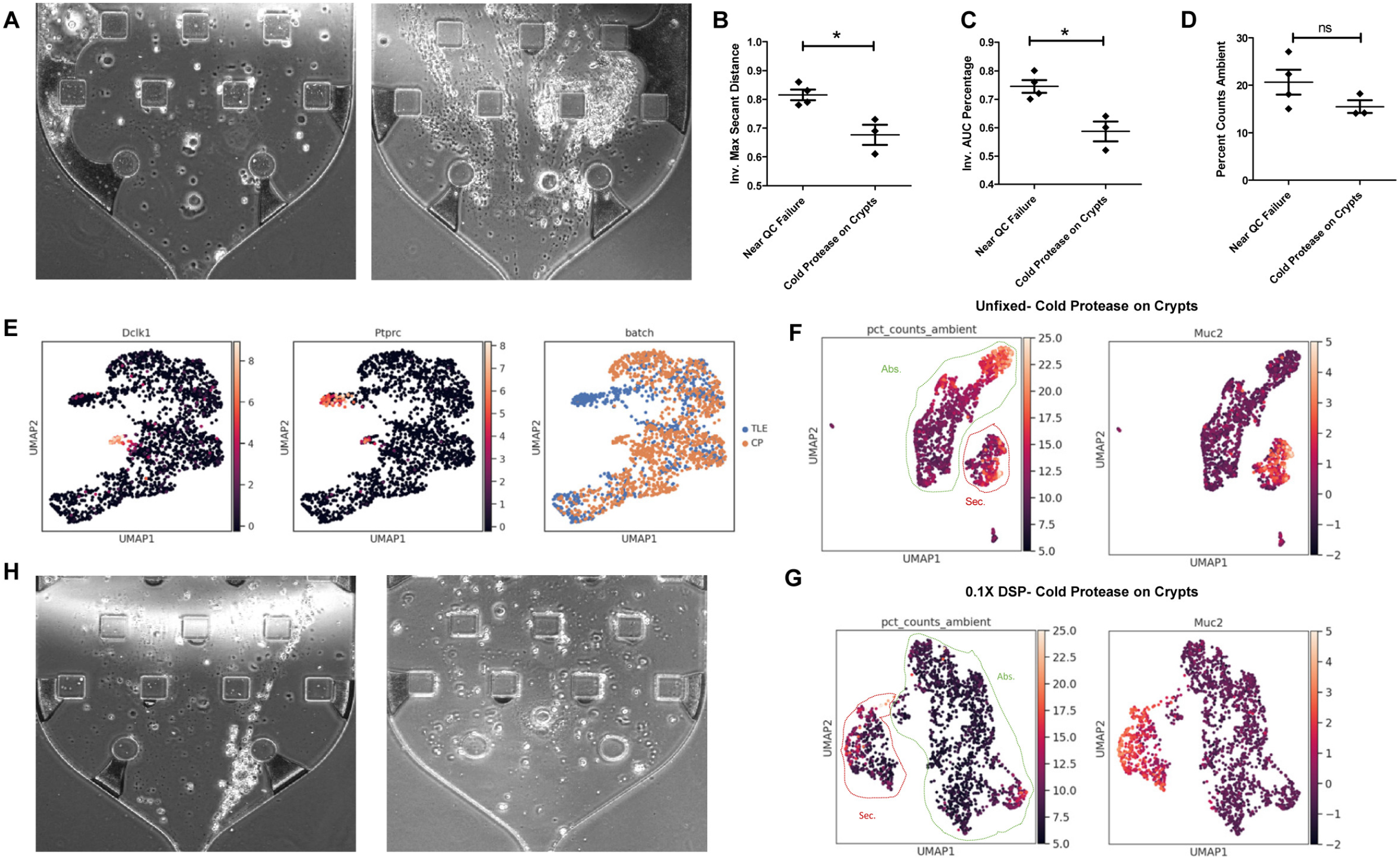
Pre-encapsulation variables affect scRNA-seq data quality and cell type diversity. **(A)** Live hopper visualization of (left) viable single cells and (right) dying cells. **(B-D)** Quantification of **(B)** inverted maximum secant distance, **(C)** inverted AUC, and **(D)** percent counts ambient comparing near QC failure runs and cold protease dissociation on crypts. Mean with SEM as error bars for n=3 or 4 samples. *p<0.05 by t-test. **(E)** UMAP embedding of filtered cells from (blue) TrypLE and (orange) cold protease datasets. Expression of *Dclk1*, a tuft cell marker, and *Ptprc*, an immune cell marker, were overlaid. **(F-G)** UMAP overlay with percent counts ambient or *Muc2* expression for **(F)** unfixed cells or **(G)** fixed cells prepared with cold protease dissociation on crypts. Secretory (red) and absorptive (green) lineages are outlined. **(H)** Live hopper visualization of (left) unfixed cells and (right) cells fixed with 0.1 X DSP.

**Table 1.** quality control metrics used on re-encasulation dissociation protocols.

|  | Contamination Metrics |  |  |  |  | Standard Metrics |  |  |  |
| --- | --- | --- | --- | --- | --- | --- | --- | --- | --- |
|  | Cell's Scaled Slope Sum | Max Secant Distance | Secant Line St.Dev. | AUC Percentage | Percent Counts Ambient | Avg. Total Number of Cells | Avg. Percent Counts Mitochondria | Avg. Total Genes Detected | Avg. Total Transcripts |
| Cold protease on Crypts | 0.62 | 0.68 | 0.402 | 0.58 | 15.52 | 1019.67 | 4.03 | 3682.52 | 12509.90 |
| MACS enzyme on Total Tissue | 0.82 | 0.78 | 0.436 | 0.7 | 27.06 | 730.00 | 9.86 | 990.96 | 1860.70 |
| Cold protease on Total Tissue | 0.78 | 0.83 | 0.450 | 0.76 | 22.34 | 295.00 | 5.06 | 1434.31 | 3220.45 |
| MACS enzyme on Crypts | 0.84 | 0.86 | 0.458 | 0.8 | 15.06 | 349.00 | 3.18 | 1989.21 | 4281.85 |
| Collagenase and DNase on Crypts | 0.8 | 0.79 | 0.436 | 0.72 | 18.14 | 249.00 | 5.83 | 2373.82 | 8155.20 |
| 0.1X DSP | 0.53 | 0.65 | 0.395 | 0.54 | 10.48 | 2049.00 | 2.05 | 3201.20 | 8209.73 |
| 1% PFA | 0.67 | 0.67 | 0.402 | 0.56 | 7.11 | 313.00 | 0.57 | 1685.39 | 4308.89 |

Because cold protease dissociation on isolated crypts resulted in the optimal balance between cell recovery and ambient contamination, we then assessed whether fixation immediately after dissociation would further improve data quality. We surmised that fixation would contain and trap all RNA within a cell and thus, will prevent ambient RNA from leaking into the surrounding buffer. We assessed two fixation strategies previously employed in scRNA-seq studies, 1% light PFA fixation used previous in combinatorial indexing (Rosenberg et al., 2018) and 0.1X dithio-bis(succinimidyl propionate) (DSP), a reversible crosslinker commonly used in pulldown studies (Attar et al., 2018). Cells fixed with 1% PFA led to QC failure, mainly due to fixation-induced degradation of RNA that disrupted library preparation **(Fig. S3K)**. However, cells fixed with 0.1X DSP led to lower contamination metrics compared to fresh tissues, again with clear delineation between secretory and absorptive cells with less ambient transcripts and non-specific *Muc2* **(Fig. 3F-G; Table 1)**. These results show that fixation can indeed contain RNA within cells and decrease ambient contamination arising from cell death during dissociation which can also be visualized in the cell hopper **(Fig. 3H)**.

### Microfluidic manipulations can affect cell death and subsequent ambient contamination in downstream data

The reduction of ambient contamination by post-dissociation fixation suggests that cells undergo continuous cell death after tissue handling. We also used live/dead cell sorting to maximize cell viability prior to cell encapsulation, which surprisingly did not improve downstream data quality (data not shown). We thus speculate that stresses encountered by cells during the microfluidic encapsulation process contribute to ambient contamination in a major way. The standard inDrops system loads cells into the encapsulation junction using channels that are 0.38 mm in diameter fed by a syringe pump. This setup results in interactions between cells and the inner surface of the channel, as well as fluid shear over relative long distances. Doubling the diameter of the loading channel to 0.76 mm marginally improved data quality based on contamination metrics **(Table 2)**. This led us to further test other strategies to reduce cell stress during microfluidic loading. The best result was obtained when cell exposure time to shear stress was minimized. To achieve this goal, an alternative cell loading setup was used to load cells directly into the encapsulation chip using a fabricated pipette tip/syringe hybrid loading system with a minimum 0.51 mm diameter to minimize cell travel time in narrow microfluidic channels **(Fig. 4A)**. The modification not only led to more viable cells as visualized on the hopper **(Fig. 4B)**, but significantly and consistently reduced contamination and increased number of captured cells **(Fig. 4C-E; Fig.S4; Table 2)**. Goblet cells were also more consistently recovered compared with standard loading **(Fig. 4F-G)**. Similar improvements were observed with total tissue dissociation in place of crypt isolation through these strategies, although data quality was lower overall, as expected **(Table S2)**. We pinpoint that the major contributor to ambient contamination and poor data quality in droplet-based scRNA-seq is the stress experienced by cells when they travel through narrow microfluidics channels.

**Table 2.** quality control metrics used on post-dissociation protocols.

| Table 2 quality control metrics used on post-dissociation protocols. |  |  |  |  |  |  |  |  |  |
| --- | --- | --- | --- | --- | --- | --- | --- | --- | --- |
|  | Contamination Metrics |  |  |  |  | Standard Metrics |  |  |  |
|  | Cell's Scaled Slope Sum | Max Secant Distance | Secant Line St.Dev. | AUC Percentage | Percent Counts Ambient | Avg. Total Number of Cells | Avg. Percent Counts Mitochondria | Avg. Total Genes Detected | Avg. Total Transcripts |
| Standard Loading Diameter = 0.38 mm | 0.62 | 0.68 | 0.402 | 0.58 | 15.52 | 1019.67 | 4.03 | 3682.52 | 12509.90 |
| Improved Loading Diameter = 0.51 mm | 0.31 | 0.46 | 0.340 | 0.34 | 3.07 | 2880.75 | 4.25 | 2803.87 | 7702.99 |
| Wide Bore Loading Diameter 0.76 mm | 0.61 | 0.67 | 0.397 | 0.58 | 11.15 | 939.00 | 3.06 | 3378.10 | 9526.15 |
| ≥ 1:1 All Cell Chip Dilution | 0.56 | 0.56 | 0.360 | 0.42 | 4.59 | 376.67 | 4.06 | 2475.67 | 8095.85 |
| 4:1 All Cell Chip Dilution | 0.82 | 0.83 | 0.450 | 0.76 | 19.24 | 156.00 | 3.54 | 1857.88 | 5408.91 |

**Fig. 4.**
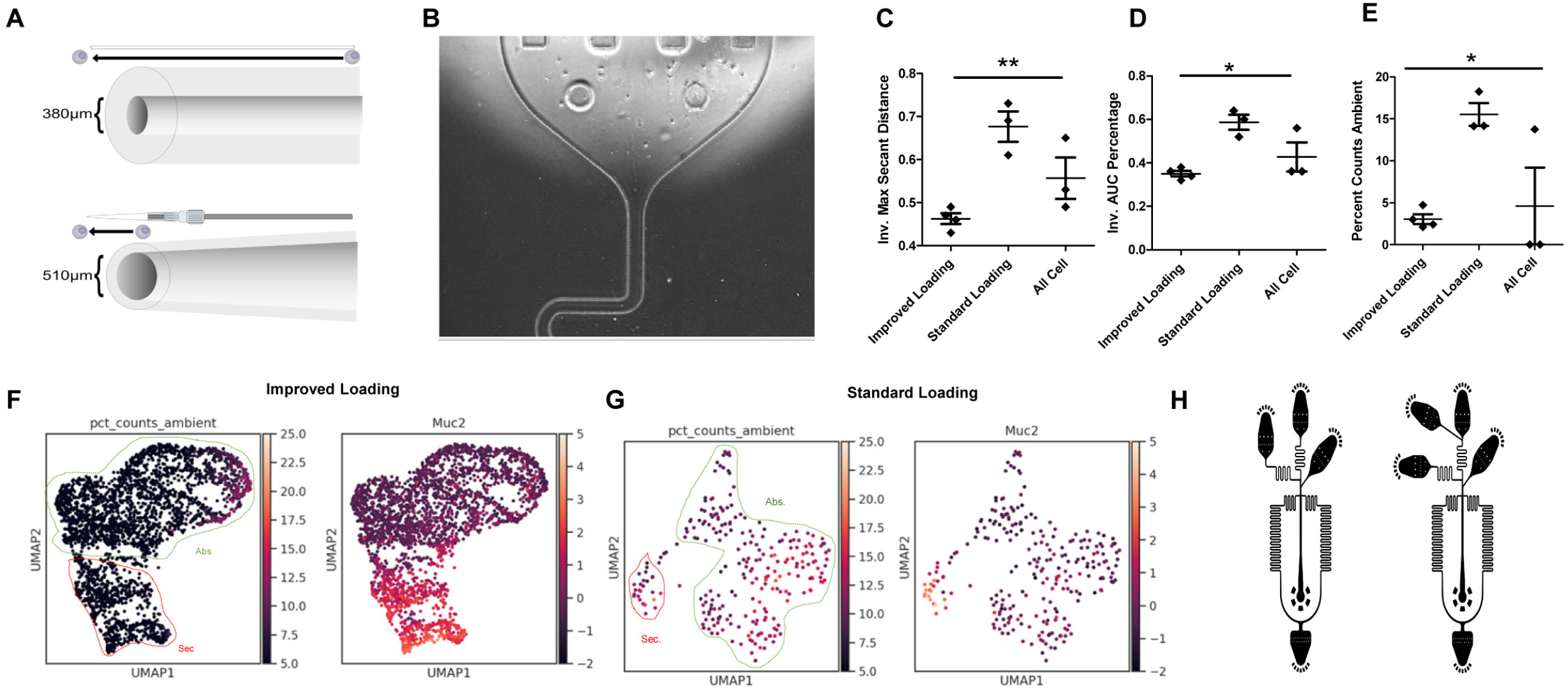
Microfluidic manipulations can affect cell death and subsequent ambient contamination in downstream data. **(A)** Schematic of standard loading (top) and improved loading (bottom). **(B)** Live hopper visualization of viable single cells from improved loading apparatus. **(C-E)** Quantification of **(C)** inverted maximum secant distance, **(D)** inverted AUC, and **(E)** percent counts ambient comparing various microfluidics manipulations. Mean with SEM as error bars for n=3 or 4 samples. *p<0.05, **p<0.01 by ANOVA. **(F-G)** UMAP overlay with percent counts ambient or *Muc2* expression for **(F)** improved loading or **(G)** standard loading. Secretory (red) and absorptive (green) lineages are outlined. **(H)** Schematic for standard inDrop chip (left), and All Cell chip (right)

We also hypothesized that ambient contamination can be reduced post-cell death via microfluidic manipulations. We utilized an alternative chip design (“All Cell”) that included another reservoir and input channel for dilution buffer, with the idea that ambient RNA in the loading buffer can be diluted out immediately before cells are encapsulated into droplets **(Fig. 4H)**. Loading the suspension in a ratio of 4:1 (cell: dilution buffer) did not improve data quality. Increasing the dilution ratio to ~1:1 resulted in lower contamination metrics **(Fig. 4C-E; Fig.S4; Table 2)**. However, due to dilution of the cell suspension prior to encapsulation, the number of cells recovered was decreased **(Fig. S4C)**. These results demonstrate that microfluidics manipulation can improve data quality pre- and post-encapsulation by tuning cell exposure to stress and diluting out ambient contamination. However, various tradeoffs, for instance, number of cells recovered, need to be acknowledged.

### Different preparations of single cells versus single nuclei yield different impact on data quality

To more universally investigate the effects of different tissue types and tissue preparation approaches, we applied our quality control strategy to the Human Tumor Atlas Pilot Project (HTAPP) dataset consisting of 40 single-cell and single-nucleus RNA-seq on 23 tumors spanning 8 cancer types generated using various protocols (Slyper et al., 2020). Samples were clustered using combinations of contamination and standard metrics and visualized as a clustered heatmap, 3D PCA plot, and UMAP **(Fig. 5A-B; Fig. S5A)**. Four clusters of sample types with differing quality were observed, with one high-quality and three lower-quality clusters. The three lower-quality clusters addressed different aspects of data quality: cluster 1 possessed lower number of captured cells/nuclei, while cluster 3 showed more ambient gene contribution **(Fig. 5A-B; Fig.S5B-J)**. Cluster 2 displayed a high number of identified cells/nuclei, even though both contamination metrics and standard metrics indicated lower data quality. We then examined different sample types and preparation conditions to elucidate factors that contribute to different data quality clusters **(Fig. 5A; Fig. S5K-O)**. The high-quality cluster was almost entirely made up of single-cell samples, whereas single-nucleus samples were abundant in the lower-quality clusters. **(Fig. 5A; Fig. S5K)**. This observation was further supported by comparing the different metrics between single-cell versus single-nucleus datasets **(Fig. 5C)**. The contamination metrics were universally higher for single-nucleus samples compared with single-cell samples, while the standard metrics were more variable. The increased number of objects captured by snRNA-seq can be attributed to a harsher and more comprehensive nuclei isolation strategy tolerated by this approach, while mitochondrial percentages were lower, being that nuclei do not contain mitochondria. Total transcript counts of snRNA-seq datasets were significantly lower than single-cell samples, since the cytoplasm was missing in such preparations. While the variable standard metrics can be attributed to various experimental factors, our contamination metrics reliably indicated that single-nucleus specimens have lower data quality than single-cell specimens. Single-nucleus isolation procedures generate nuclei containing adhered ribosomes and RNA that can easily shed into the loading buffer to create ambient contamination.

**Fig. 5.**
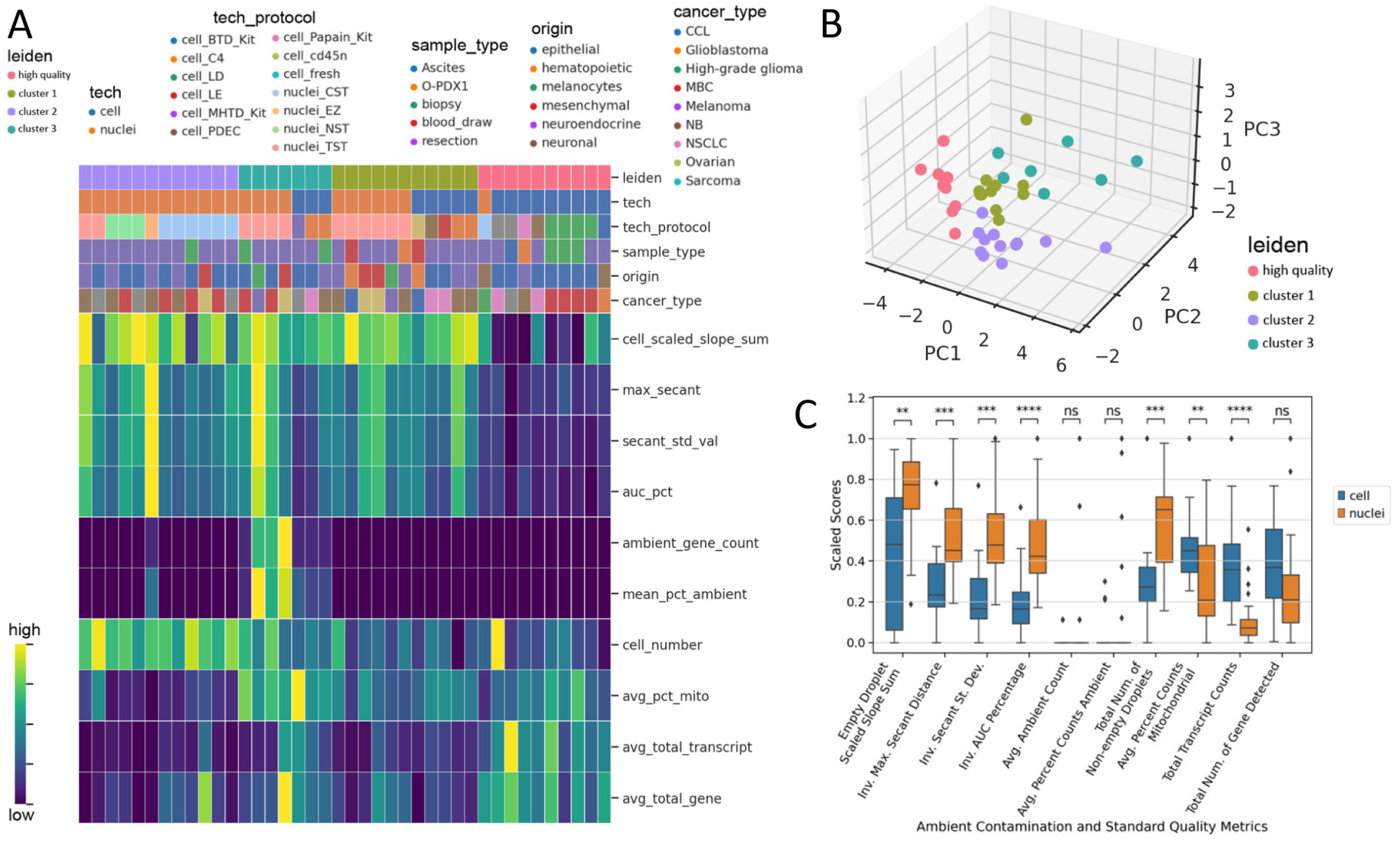
Ambient contamination and quality control metrics reveals impact of intrinsic and extrinsic factors on data quality. **(A)** Heatmap of ambient contamination and standard QC metric scores with HTAPP datasets as columns grouped by Leiden clusters. Metrics are shown as rows. The Leiden cluster labels and labels of isolation technique, technique x protocol, sample type, tissue origin, and cancer type are shown as color bars above the heatmap. Metric scores are normalized between 0 and 1 for each row for visualization. **(B)** Three-dimensional scatter plot of the first 3 principal components of the ambient contamination and standard QC metric score matrix colored by Leiden cluster labels. **(C)** Boxplot comparing the metric scores between single-cell and single-nucleus sequenced samples. Two-sided Mann-Whitney-Wilcoxon test performed between single-cell and single-nuclei groups. **p<0.01, ***p<0.001,****p<0.0001.

Further inspection indicated that certain techniques such as single-cell Liberase and DNase (LD) and CD45+ depletion (CD45n) were consistently yielding higher-quality data, whereas other protocols such as single-cell C4 (Collagenase 4 and DNase I) resulted in more contaminated datasets **(Fig. 5A; Fig. S5L)**. There were also differences in snRNA-seq approaches. For example, Tween with salts and Tris (TST) was in cluster 1 and 3 with more mitochondrial and ambient contribution, whereas Nonidet P40 and CHAPS with salts and Tris (NST and CST) generally presented in cluster 2 with higher number of nuclei identified. However, cancer type, tissue origin, or tissue collection procedures scattered randomly amongst clusters, indicating no concordance with data quality **(Fig. 5A; Fig. S5M-O)**. These results indicate that the cell/nuclei preparation protocol was the major factor impacting data quality.

## Discussion

The democratization of droplet-based scRNA-seq technology has led to its widespread application in understanding tissue biology and heterogeneity. Large consortia have been established for human tissue profiling using scRNA-seq as a central approach (HuBMAP Consortium, 2019; Rozenblatt-Rosen et al., 2020). However, several technical artifacts, such as ambient contamination, have not received sufficient attention. The contamination problem is particular insidious for healthy tissues compared to cancer tissues, as differentiated cells, especially fragile secretory cells, are more prone cell death and stress. The most prevalent mode for quality control of a scRNA-seq dataset is the ability to distinguish cell populations via marker genes in some reduced dimension space. Other standard metrics such as feature counts, transcript counts, and cells recovered do not directly reveal ambient contamination. While canonical cell types may still be identified by highly expressed marker genes in a dataset heavily contaminated by ambient RNA, downstream analysis regarding gene programs, states, and pathways will be significantly confounded. For instance, “mixed lineage” cells may just be artifacts of ambient contamination and not of real biology. We also speculate that a large portion of batch effect may arise from ambient contamination whose composition is random for every set of experiment. Furthermore, ambient contamination may even prevent real cells from being effectively distinguished from empty droplets, resulting in significant manual efforts in filtering post-QC. Given the widespread use of scRNA-seq, we suspect that there are potentially many published datasets fraught with ambient contamination to different degrees, and end users of publicly available data would benefit from having tools to assess the severity of this problem. We provide quantitative metrics using post-alignment counts data prior to and post filtering that reveal ambient contamination in scRNA-seq datasets. Our contamination metrics essentially capture ability to identify signal (biological transcripts) from noise (ambient transcripts) within a scRNA-seq experiment. Aside from ambient contamination that contribute to noise, our metrics can also identify experiments with excessively low transcript counts that approach baseline signals. Low number of cells does not necessarily equate to poor quality as encapsulation of a small number of highly viable cells can still result in high signals. Nevertheless, cell number can affect data quality if the starting material becomes so minute that the ability to amplify cDNA and prepare sequencing-amenable libraries is affected.

Several computational packages have been developed to address ambient contamination post-hoc. EmptyDrops, dropkick, and DropletQC leverages the ambient contamination profile for automatically identifying cells from empty droplets (Heiser et al., 2021; Lun et al., 2019; Muskovic and Powell, 2021). DecontX, SoupX, and CellBender are tools for factoring out the contribution of ambient RNA in the resultant counts data matrix (Fleming et al., 2022; Yang et al., 2020; Young and Behjati, 2020). However, we surmise that addressing the problem experimentally on the front end will be the best strategy. We established that cell death is the main culprit of ambient contamination from the presence of leaking chromatin in the microfluidic encapsulation chip. Leaked DNA can lead to a cascade of cell death events, as nucleic acid material can act as danger signals for TLR9 and cGAS (Paludan et al., 2019). We pursued strategies to minimize cell stress in a scRNA-seq experiment. We validated that mechanical stresses of tissue handling and length of high temperature dissociation affect cell quality, although failure at these steps leads to QC failure that is easy to detect with standard metrics and data analysis. The main contributor to ambient RNA lies in post-dissociation microfluidics encapsulation, specifically the stresses experienced by cells as a function of length of distance travelled and channel diameter. The concentration of ambient contamination in the loading buffer can also be manipulated at encapsulation. The degree of contamination also varies by protocol, with single nuclei sequencing possessing more severe ambient contamination. While microfluidics parameters cannot be easily altered in commercial systems, there are now multiple platforms with different encapsulation strategies that can be selected for different applications using the knowledge derived from this report.

A common misconception in the field is the efficacy of snRNA-seq in dealing with cell death and ambient RNA. Nuclei are thought to be more hardy, smaller in size, and more resistant to mechanical damage. We did not see an improvement in data quality in snRNA-seq data. Isolated nuclei have adhered cytoplasmic RNA that are dislodged into the loading buffer, contributing to ambient contamination. There is support for this adhered RNA in the literature, as the nuclei-attached ribosomes can be visualized (Slyper et al., 2020) and snRNA-seq data have been used for RNA Velocity that utilizes spliced RNA (Marsh and Blelloch, 2020). Nuclei also have smaller RNA pools that result in lower capture efficiencies and biological signals, which can be overwhelmed by ambient RNA. While snRNA-seq makes single-cell analysis possible for frozen and difficult-to-dissociate tissues (May-Zhang et al., 2021), end users should balance these factors when deciding whether to use cells or nuclei for their single-cell analysis.

## STAR METHODS

Detailed methods are provided in the online version of this paper and include the following:

- KEY RESOURCES TABLE
- RESOURCE AVALIABILITY
  - Lead contact
  - Materials availability
  - Data and code availability
- EXPERIMENTAL MODEL AND SUBJECT DETAILS
- METHOD DETAILS
  - A. Droplet simulation
  - B. Total tissue dissociation
  - C. Crypt isolation
  - D. Single-cell dissociation
  - E. Fixation
  - F. inDrops Encapsulation
  - G. Dilution microfluidics encapsulation chip
  - H. Quality Control Metrics
- STATISTICAL ANALYSIS

## SUPPLEMENTAL INFORMATION

Supplemental information can be found online at

## INCLUSION AND DIVERSITY

One or more of the authors of this paper self-identifies as an underrepresented ethnic minority in science. One or more of the authors of this paper received support from a program designed to increase minority representation in science.

## ACKNOWLEDGEMENTS

The authors wish to thank other contributing investigators, including Marisol Ramirez-Solano, Changqing Zhang, Thomas Wise, Lori Coburn, Keith Wilson, and Ian Hurford, as well as the Vanderbilt Epithelial Biology Center for insightful discussions. This study was supported by U2CCA233291, P50CA236733, R01DK103831, and U54CA274367 from the NIH, G-1903-03793 from The Leona M. and Harry B. Helmsley Charitable Trust (to KSL), T32LM012412 (in support of BC), CA190172 from the DoD, R37CA244970 and R01CA272687 from the NIH, the AGA-R. Robert & Sally Funderburg Research Award in Gastric Cancer (to EC). We would like to acknowledge the VANTAGE core supported by P30CA06848.

## AUTHOR CONTRIBUTIONS

Conceptualization (DA, ZC, AJS, KSL); Data curation (DA, AJS); Formal analysis (DA, ZC, KSL); Investigation (DA, ZC, CJHB, EC, JDC, QL, KSL); Methodology (DA, ZC, AJS, CNH, ANS, MJB, YY, BC, YX, KSL); Project administration (KSL); Resources (KSL); Software (ZC); Supervision (KSL); Validation (DA, ZC, QL); Visualization (DA, ZC, KSL); Writing-original draft (DA, ZC, KSL); Writing-reviewing & editing (DA, ZC, CJHB, EC, JDC, QL, KSL)

## KEY RESOURCES TABLE

### RESOURCE AVAILABILITY

#### Lead contact

Further information and requests for resources and reagents should be directed to and will be fulfilled by the lead contact, Ken Lau

#### Materials availability

This study did not generate new unique reagents.

#### Data and code availability

The contamination metrics pipeline developed for this study can be found at (https://github.com/Ken-Lau-Lab/AmbientContaminationMetrics.git)

### EXPERIMENTAL MODEL AND SUBJECT DETAILS

Animal experiments were performed under protocols approved by the Vanderbilt University Animal Care and Use Committee and in accordance with NIH guidelines. Wild-type mice (C57BL/6) of both sexes were euthanized in an approved fashion prior to dissection and tissue harvesting.

#### A. Droplet Simulation

We used CellBender (Fleming et al., 2022) to simulate representative datasets of different quality. We generated synthetic datasets by randomizing the number of real cells, number of droplets, number of genes/transcripts, over distributions centered on 2000, 12000, and 5000 respectively. The ambient transcript levels were set between 100 to 4000 to simulate a range of ambient contamination levels. n=1000 simulations were performed for each ambient level.

#### B. Total Tissue Dissociation

Previously established protocols for total tissue dissociation were followed (https://dx.doi.org/10.17504/protocols.io.busfnwbn). Briefly, tissues were minced into 1-2mm pieces using a scalpel followed by enzymatic incubation for dissociation.

#### C. Crypt Isolation

Colonic crypt isolation was performed as previously documented (Liu et al., 2018). Briefly, isolated colonic tissues were chelated in buffer consisting of 3 mM EDTA (Corning) and 0.5mM DTT (Teknova) in 1X Dulbecco’s Buffered Saline (DPBS) for 1 hour and 15 minutes rotating at 4 *°C*. The tissue was then transferred to 1X DPBS and shaken rigorously for 2 minutes to separate the colonic epithelium from the tissue. After transfer of crypts to a new tube, shaking was repeated 3X to collect remaining crypts.

#### D. Single-cell Dissociation

For colon, minced tissue or crypts were dissociated either using cold protease, MACS enzyme, or a DNase1/collagenase enzymatic cocktail. For testing the enzymatic cocktail found in MACs Mouse Tumor dissociation kit (Miltenyi Biotech), tissues were incubated 20 minutes with gentle orbital shaking (~200RPM) at 37*°C*. For dissociation with the cold protease cocktail, tissues were incubated with cold protease (Sigma-Aldrich) (5mg/ml) and DNase (Sigma-Aldrich) (2.5mg/ml) on a rotator (~8 rpm) for 25 minutes at 4*°C*. For dissociation using Collagenase (2mg/ml) with DNase (2.5mg/ml), tissues were incubated for 20 minutes at 37*°C* static with trituration at 10-minute intervals (Banerjee et al., 2020). After enzymatic incubations, gentle pipetting with a wide bore p1000 pipette was used to mechanically dissociate tissues, resulting in visibly turbid cell suspensions. After dissociation, the digestion enzyme mixtures were quenched with 2% FBS, and the suspensions were passed through a 70μm filter (Pluriselect) to generate single cells. A series of washes were performed to obtain an optimal single-cell suspension to minimize debris.

For stomach corpus, the mucosa was scrapped off using cell scrapers and incubated in the pre-warmed digestion buffer (1mg/mL Collagenase, 2.5mg/ml DNAse) on a 37 °C shaker, at 220 rpm for 30 minutes. After quenching and filtering, the glands were pelleted at 300 g for 5 minutes and dissociated further in TrypLE (Gibco) and Y-27632 at 37 °C for 5 min, and was quenched and spun down at 500g for an additional 5 minutes thereafter prior to encapsulation.

#### E. Fixation

Single-cell suspensions were fixed with 1% paraformaldehyde (PFA) (Thermo Scientific) or 0.1X DSP (Thermo Scientific). DSP was solubilized in 100% DMSO (Sigma-Aldrich) at a final concentration of 25 mg/mL to form a “25×” stock. 1X DSP solution was prepared and filtered as previously documented (Attar et al., 2018). This solution was then added dropwise to single-cell suspensions to achieve the final concentration. Samples were incubated on a rotating platform for 30 minutes at room temperature. Residual DSP was then quenched with Tris-HCl added to a final concentration of 20 mM. Reverse cross-linking was conducted by reducing the disulfide bonds of the DSP fixative using the 10mM DTT present in the standard inDrops RT/Lysis buffer.

#### F. inDrops Encapsulation

For standard inDrops, the protocol outlined previously was followed (Zilionis et al., 2017). For library preparation, the TruDrop library structure and preparation were adopted (Southard-Smith et al., 2020). Alignment of reads and barcode deconvolution to generate count matrices was performed using the DropEst pipeline (Petukhov et al., 2018). Modifications of inDrops were made as documented below and described in (Chen et al., 2021a; Simmons and Lau, 2022). Standard loading was performed with assemblies made using 0.38 mm inner diameter tubing 20 cm in length (Scientific Commodities) and fed with a syringe pump. Wide bore loading was performed similarly but with 0.76 mm inner diameter tubing (Scientific Commodities). For enhanced loading, luer lock connectors (Qosina) were spray coated with primer (Loctite) and allowed to dry for 10 minutes in a fume hood. After drying, connectors were glued to pipette tips (Biotix) using UV curing glue (Loctite) and cured using a UV Crosslinker (Stratalinker). Cells were then loaded into the tip assembly, locked using male to female luer adaptors (Idex) to a syringe assembly made using a 30 cm length of tubing (Cole-Parmer), primed with mineral oil (Sigma-Aldrich) colored with oil red (Alfa Aesar), and connected to a syringe pump. Red mineral oil then acted as a void volume to push cells directly into the inDrops encapsulation chip.

#### G. Dilution Microfluidic Encapsulation Chip

Encapsulations that incorporate dilutions were performed using the “All Cell” chip design as shown in **Fig. 4H**. The chip design was nearly identical to the standard inDrops chip but featured an additional inlet connection to the cell channel for diluting cells immediately prior to entering the encapsulation junction. A syringe pump (New Era Pump Systems) was used to push DPBS into the chip prior to droplet partitioning through this additional channel. Flow rates for cell and dilution buffers were adjusted to total a rate matching that of the reverse transcriptase, such that enzyme and buffer conditions in the final droplets were kept constant.

#### H. Quality Control Metrics

##### Data processing

To apply the ambient contamination metrics on a dataset, the first step was to read-in raw gene count data and scale the dataset barcode number relative to the expected real cell number to enable comparison between samples with different numbers of encapsulated cells and empty droplets. The starting point was an unfiltered count matrix that can be of various formats (h5ad, mtx, etc.). An inflection curve was computed using the find_inflection() function from QCPipe.qc module (Chen et al., 2021b) where a cumulative sum curve of total transcript counts vs barcodes ranked by their transcript counts would be drawn, and the first inflection point of the curve would be used as an estimated real cell number for the sample. This estimation was based on the rationale that, when ambient RNA contamination is not comparable to the true biological transcript counts, droplets capturing real cells contribute to distinctly larger increments in the cumulative count value. In contrast, empty droplets contribute less, so the first inflection point could be a position to approximate captured real cells versus empty droplets. Alternatively, an estimated real cell number could be manually entered in our function as an argument if an expected real cell number for the sample was known. After determining the estimated real cell number, barcodes would be sorted based on their total transcript counts; a threshold would be set as a multiple of the estimated cell number to retain only the high transcript count barcodes beyond the threshold. Through observations of samples used in this study, the multiple was set to 4 as default. This data-processing step could be performed using the cut_off_h5ad() or cut_off_from_dropset() functions from our data_processing python module.

##### Geometric quantification of ambient contamination from the cumulative transcript count curve

Our ambient contamination metric calculation integrated geometric quantifications of the cumulative transcription count curve (**Fig. 1 A-B**). As the total barcode number was set to a multiple of the estimated real cell number, the curve of a high-quality dataset was expected to raise with steep slope in the first portion 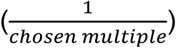, then turn to a relatively flat slope. This shape could cause the curve to deviate notably away from the diagonal linking the final cumulative count and the origin initially (**Fig. 1A**) then gradually get close to the diagonal. However, in low quality cases where ambient RNA molecules keep contributing to high increments of the cumulative count throughout the dataset, the cumulative count curve’s slope would have a low variance, and the curve would not deviate far from the diagonal substantially (**Fig. 1B**). Therefore, the magnitude of the curve’s deviation from the diagonal line and the variance within the distances between the curve and the diagonal line are indicators of data quality. We computed the vertical distances between the cumulative sum curve and the diagonal for each barcode, which we defined as secant lines whose maximal value and standard deviation were then calculated to inform data quality. To establish quantitative indicators positively correlated with ambient levels, we inverted the maximal secant distance and the standard deviation by the subtractions:

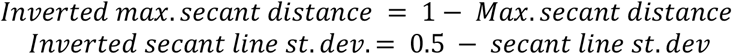

The rationale was that the cumulative transcript count curve was normalized to a range of 0 to 1, so all secant lines’ length should fall between this range. In extreme cases of secant lines with a maximal distance close to one, a minimal distance close to 0, and a minimal sample size (eg. =3), the standard deviation did not go beyond 0.5 and will always be positive. We used the inverted standard deviation and maximal value of the secant lines as two metrics.

In addition to the secant line distances, as we have described previously, a high-quality dataset has a cumulative count curve resembling a rectangular hyperbola with a sharp incline and then flattening of the curve. In contrast, the lack of deviation from the diagonal line for low quality datasets makes the curve resemble a straight line. We quantified the shape differences by first computing the minimal rectangle area circumscribing the cumulative count curve:

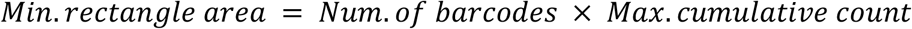

We then computed the area under the cumulative count curve (AUC) using sklearn.metrics.auc() function. Taking the ratio between the area under the cumulative count curve and the area of the minimal rectangle would give a percentage value.

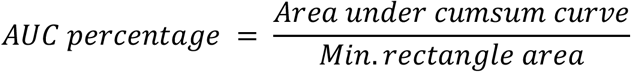

We inverted the AUC percentage by quantify the area above the curve within the minimal rectangle:

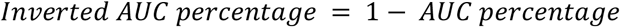

A higher inverted AUC percentage value indicates the closeness of the area to the triangle formed from x,y axes and the diagonal, thus high contamination. Therefore, Inverted AUC percentage was used as one of our ambient contamination metrics. Steps to compute metrics before inversion were encapsulated in our plot_quality_score.plot_secant_line() function. Alternatively, numerical results alone could be computed from our calculation.secant_metrics() function.

##### Statistical quantification of ambient contamination from the distribution of the slope of the cumulative transcript count curve

As described earlier, high quality datasets have the pattern of a sharp incline followed by flattening of their cumulative transcript count curves, whereas low quality datasets have curves with small change in slopes. The slope difference can be a continuous value informing the quality in a quantitative way. We therefore inspected the slope distribution by generating a histogram on the slopes at each barcode for a sample (**Fig. S1I-J**) using matplotlib.axes.Axes.hist() function, fixing the parameter of number of bins at 100 for consistency. We expected to see a bimodal distribution with a peak contributed by low-slope barcodes followed by a peak contributed by high-slope barcodes, and a higher density at high slope region for high-quality datasets than low-quality datasets was expected. However, even though the two modes could be observed on histograms of high-quality datasets (**Fig. S1I**), it was hard to compare the density values because most droplets in the dataset were expected to be empty due to the way we cut-off the dataset as described above, rendering a notably heavier mass contributed by low-slope barcodes than high-slope barcodes.

To emphasize the density contribution of the high slope barcodes, we transformed the distribution to scale up the weights of high-slope barcodes. We performed the transformation by getting each histogram bin’s mean slope value as the x-value for the transformed plot and each bin’s frequency value multiplied by the bin’s mean slope as the y-values for the transformed plot (**Fig. 2C-D**). In this way, the contribution of distribution density from high-slope barcodes were scaled up based on their slope value, and we were able to quantify the scaled density contributed by these high-slope barcodes as one metric.

To determine slopes that are likely contributed by real cells rather than empty droplets, we set a threshold to have a binary assignment of a barcode to be either real cell or empty droplet. The threshold was calculated as:

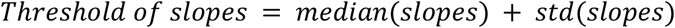

Barcodes with slopes higher than the threshold will be identified as real cells. Since we wanted to include only distinguishably high-slope barcodes as real cells, in using the median value, the threshold was less skewed by the high density of low-slope barcodes. In adding one standard deviation, we accounted for the fact that real cells are expected to be a small portion of the dataset, so a quantity of the dataset’s internal variation added to the median could further refine the threshold to include potential real cell barcodes with high slope contribution.

We transformed the scaled slope distribution so that the area under the curve summed up to one. Summing the y-values contributed by barcodes beyond or below the threshold gave us the scaled slope contribution from potential real cells and empty droplet respectively. We used the scaled slope sum from the low-slope region (potential empty droplet) as another metric whose value would increase with increased ambient level. Histograms were be generated by our plot_quality_score.plot_slope() function, whose return value could be passed to our plot_quality_score. plot_freq_weighted_slope() to generate the scaled slope distribution plot. Alternatively, numerical results alone could be computed from the calculation.freq_slope_area_ratio() function.

##### Ambient gene quantification

Ambient genes were defined as genes that have a dropout rate of less than 2% in this study. Dropout rate of each gene was calculated, and the dropout rate vs ranked gene plot was generated as described in (Heiser et al., 2021). Percentage counts of ambient genes were computed with scanpy.pp.calulate_qc_metrics() function, with the specific argument ‘qc_vars = [“ambient”]’, where ‘ambient’ was an anndata object’s obs variable composed of a list of boolean labels identifying the ambient genes among all genes. Histograms of the distribution of percentage counts of ambient genes were made with seaborn.histplot() function.

##### Standard quality control metrics calculation

We followed the steps of (Chen et al., 2021b) to filter barcodes based on dropkick scores, standard QC information and biological markers. Total transcript counts, total counts of gene detected and percentage counts of mitochondrial genes were calculated with scanpy.pp.calulate_qc_metrics() function on datasets after filtering and were used as the standard QC metrics in our study.

##### UMAP Visualization

For single-cell RNA-seq data, we normalized raw count data by median library size, log-like transformed with Arcsinh, and Z-score standardized per gene using scanpy and numpy functions. For the metrics scores matrix, we performed the Arcsinh transformation and Z-score standardization with scapy and numpy functions. UMAP coordinates were calculated after PCA and KNN clustering on the matrices as described in (Chen et al., 2021b) and UMAPs were visualized with scanpy.pl.umap() function.

##### Heatmap and 3-D Scatter Plot Visualization

The heatmap was generated using seaborn.clustermap() function with the input of matrix of the 6 ambient contamination metric and the 4 standard QC metric scores as rows and the HTAPP samples as columns. The matrix was sorted by samples’ Leiden cluster labels and then by sequencing technique & protocol labels before making the heatmap;‘col_cluster’ and ‘row_cluster’ were set to false as the function’s input. Metadata of sequencing techniques, sequencing technniques & protocols, sample types, cell origins and cancer types were input as a list of mapped colors as the col_colors argument to the function. The 3-D scatter plot was generated with matplotlib.Axes.scatter() function; input x,y,z coordinates were the 3 principal components of the HTAPP sample’s ambient contamination and standard QC metric scores after Principal Component Analysis (PCA). PCA was performed using scanpy.tl.pca() function with svd_solver = ‘arpack’ argument after the metric score matrix was archsinh transformed and scaled to unit variance and 0 mean using numpy and scanpy functions.

### QUANTIFICATION AND STATISTICAL ANALYSIS

p-values from two-sided, unpaired t-test and One-way ANOVA, Tukey test. P-values below 0.05 are considered statistically significant.

## Supplementary Fig. Captions

**Fig. S1. Distribution of simulation parameters and simulated data related to Fig. 1. (A-B)** Distribution of the number of **(A)** droplets and **(B)** real cells per dataset for 1000 datasets simulated by CellBender. **(C)** Distribution of the number of biological transcripts UMI counts per cells for a dataset with ~2000 cells simulated by CellBender. **(D)** Distribution of the number of ambient UMI counts per droplets for various datasets at different ambient levels. 120,000 droplets per dataset simulated by CellBender. **(E-F)** UMI count versus log ranked barcodes curve for datasets simulated with **(E)** low ambient level and **(F)** high ambient level. **(G-H)** Scaled cumulative total transcript counts over ranked barcodes by total transcript counts for datasets simulated with **(G)** low ambient level and **(H)** high ambient level. Slopes (dashed pink lines) distinguishing real cells and empty droplets can be observed in **(G)**, but not in **(H)**. **(I-J)** Distribution of the slopes of the for curves shown in **(E-F)**, respectively. **(K)** Number of ambient genes over ambient levels for simulations. Line plots shown as mean ± stdev of n=1000 replicates for each ambient level.

**Fig. S2. Quality control metrics that quantify ambient contamination in experimental datasets related to Fig. 2. (A-H)** Ambient contamination plots and metrics formatted similarly to **Fig. 2** of K562 sample 2 **(A-D)** and K562 sample 3 **(E-H)**. **(I-M)** UMAPs of filtered cells overlaid with mitochondrial count percentage for **(I)** K562 sample 1, **(J)** K562 sample 2, **(K)** K562 sample 3, **(L)** mouse gastric corpus, and **(M)** mouse colonic crypts. Absorptive and secretory cell clusters are delineated by dashed green and red lines, respectively. **(N)** UMAP of the colonic crypts dataset overlaid with *Muc2* gene expression level.

**Fig. S3. Quantitative assessment pre-encapsulation factors that affect data quality related to Fig. 3. (A-D)** Ambient contamination plots formatted similarly to **Fig. 1** derived from colonic datasets generated using various dissociation protocols, showing near QC failure. **(E-J)** Quantification of **(E-F)** contamination and **(G-J)** standard metrics comparing near QC failure runs and cold protease dissociation on crypts. Mean with SEM as error bars for n=3 or 4 samples. *p<0.05 by t-test. **(E)** UMAP visualization colored according to gene expression for cell type markers for 1% PFA fixation colonic crypt scRNA-seq dataset.

**Fig. S4. Evaluation of post-dissociation factors in affecting data quality related to Fig. 4 (A-F)** Quantification of **(A-B)** contamination and **(C-F)** standard metrics comparing various microfluidics manipulations. Mean with SEM as error bars for n=3 or 4 samples. *p<0.05, **p<0.01 by ANOVA. **(G)** Table of Tukey’s post-test values calculated for each metric.

**Fig. S5. Ambient contamination, quality control metrics and sample metadata visualization related to Fig. 5**

UMAPs of ambient contamination and standard QC metric scores overlaid with **(A)** cell’s scaled slope sum, **(B)** maximal secant distance, **(C)** secant line standard deviation, **(D)** AUC percentage, **(E)** average ambient gene counts, **(F)** average percentage counts of ambient genes, **(G)** total number of non-empty droplets, **(H)** average percentage counts of mitochondrial genes, **(I)** average total transcript counts, **(J)** average total gene detected, **(K)** isolation technique, **(L)** technique x protocol, **(M)** sample type, **(N)** tissue origin, **(O)** cancer type. The high-quality cluster is circled in red dash line, and low-quality clusters are circled in blue dash lines.

