## Supplementary figures and images for "A contamination focused approach for optimizing the single-cell RNA-seq experiment"

### Supplemental Figure 1

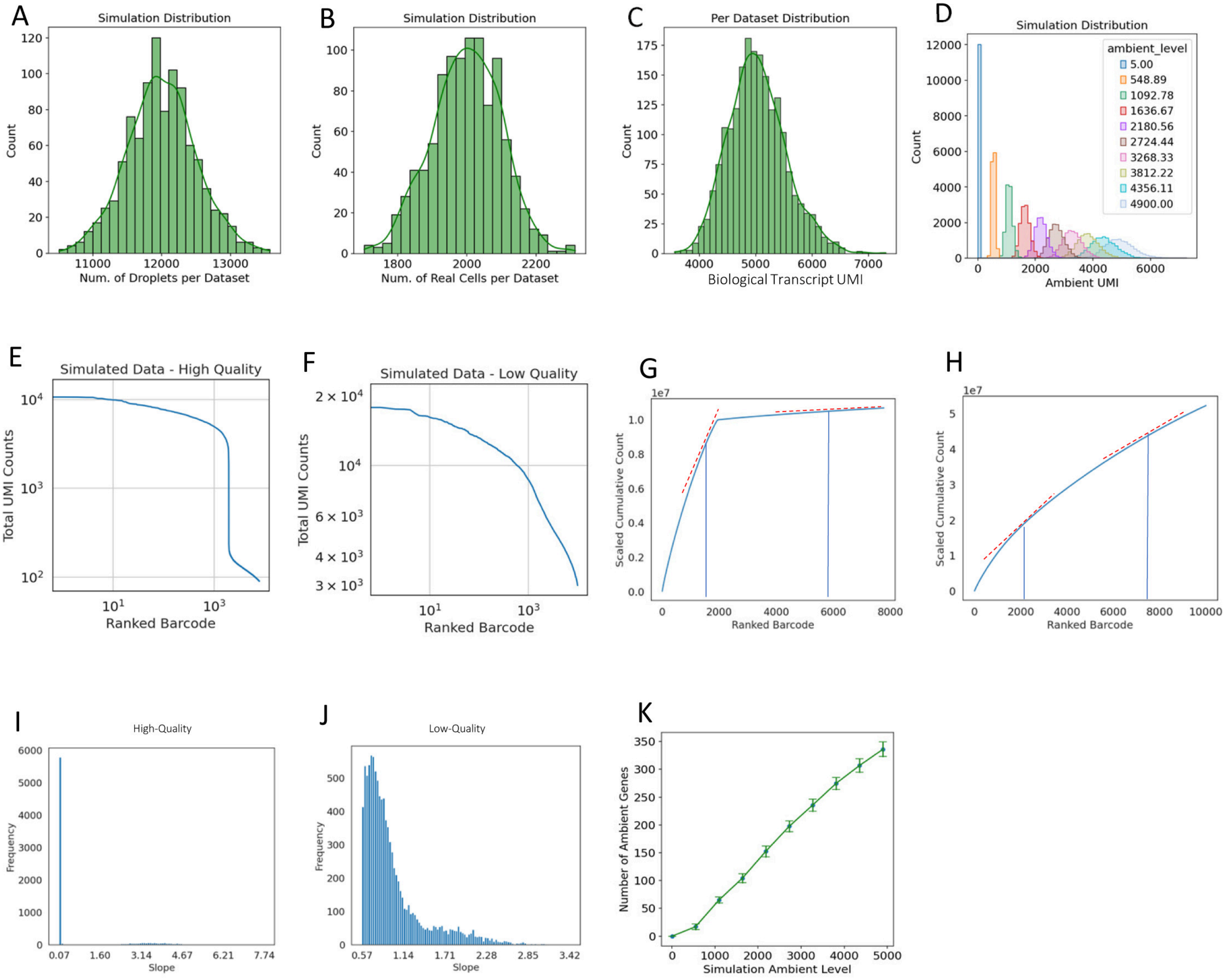

### Supplemental Figure 2

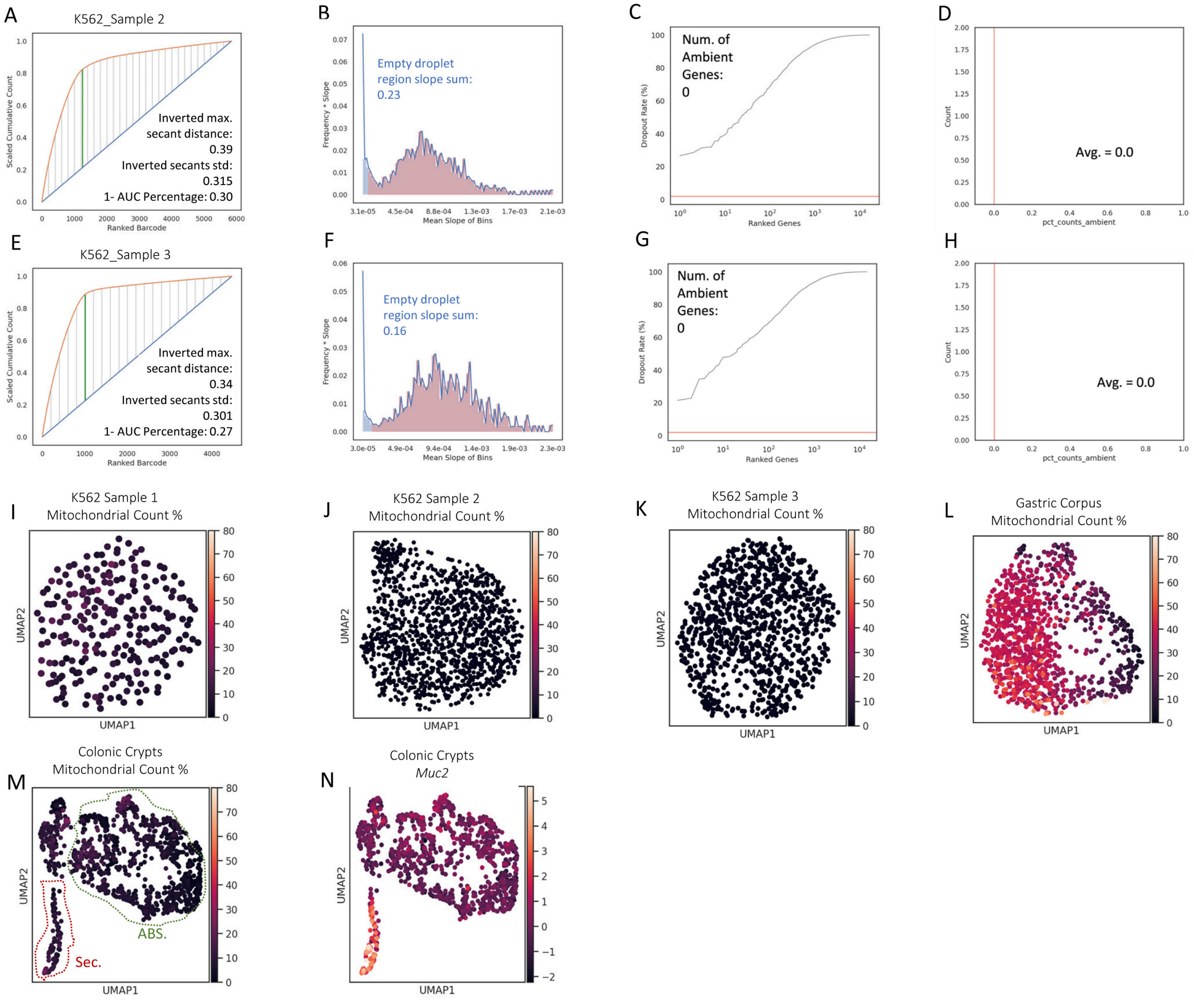

### Supplemental Figure 3

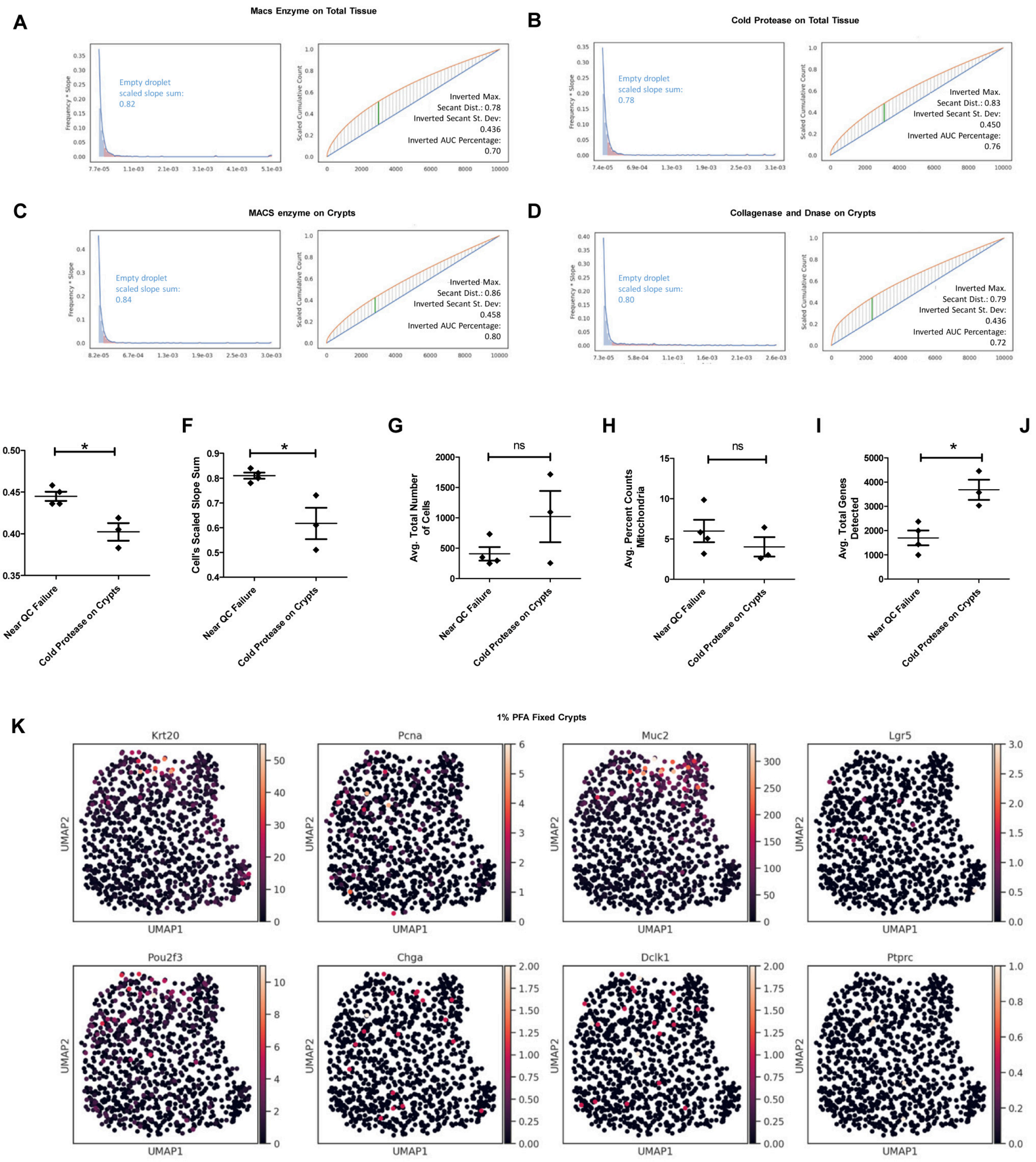

### Supplemental Figure 5

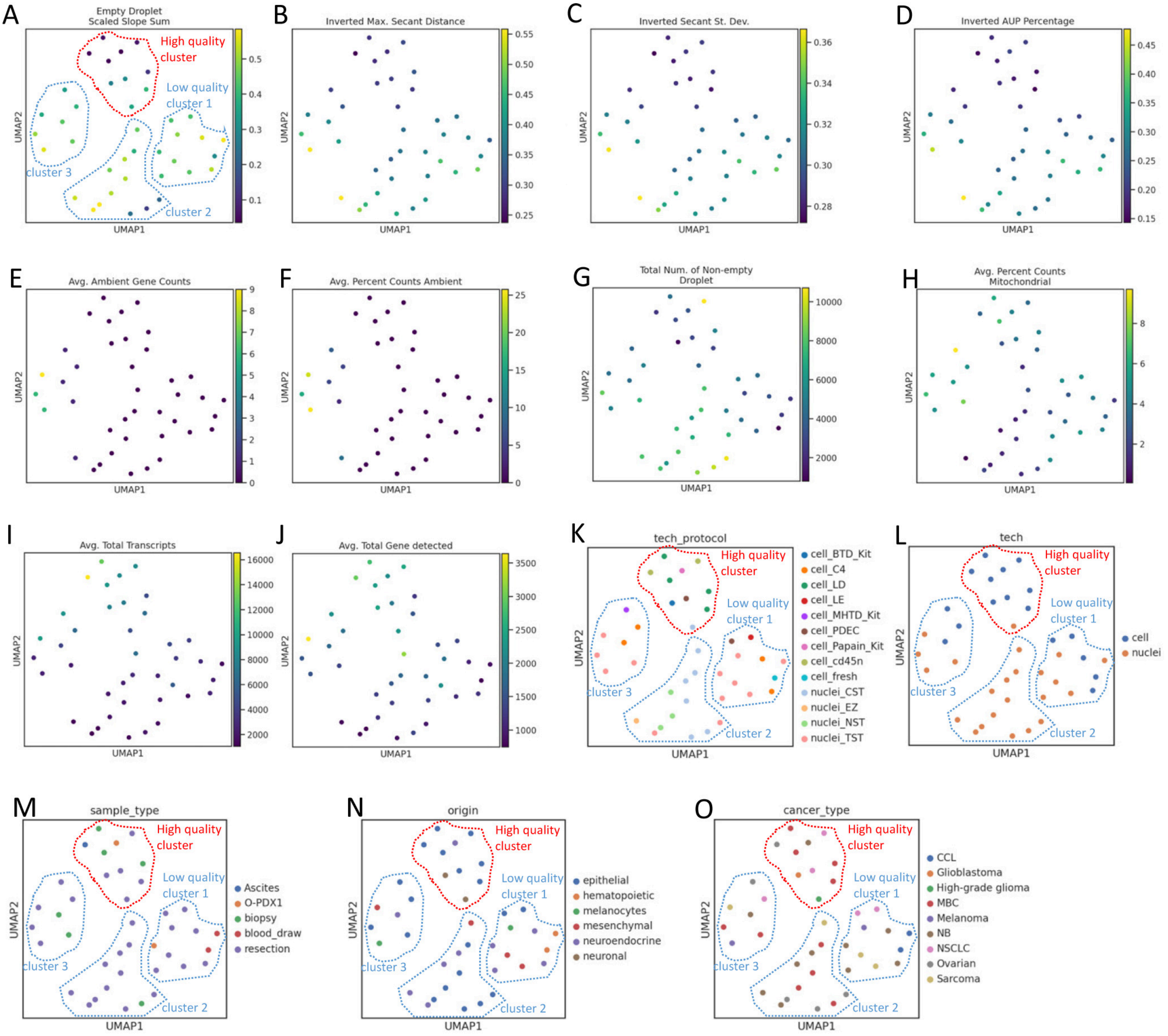
