## Supplemental Figure 4 for "A contamination focused approach for optimizing the single-cell RNA-seq experiment"

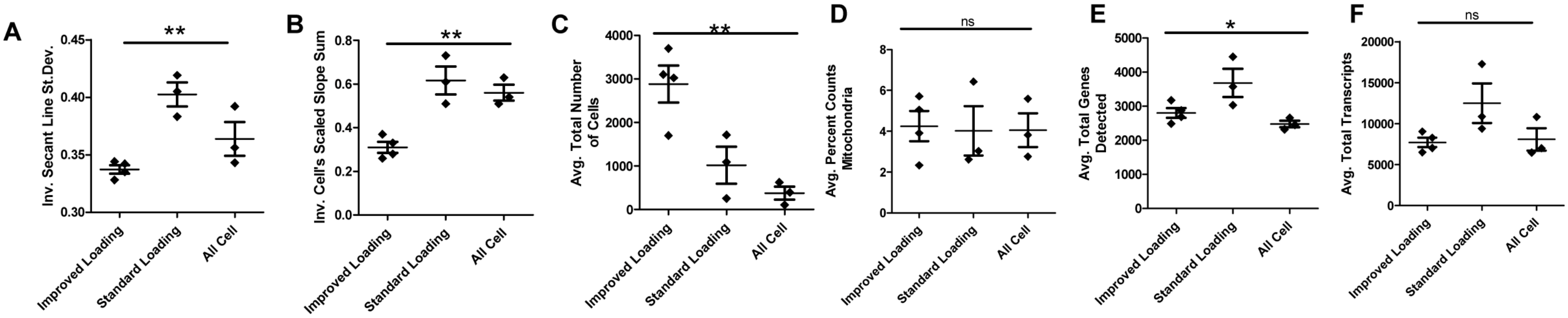


G

| Inv. Max Secant Distance | | | | | |
|---|---|---|---|---|---|
| Tukey's Multiple Comparison Test | Mean Diff. | q | Significant? P < 0.05? | Summary | 95% CI of diff |
| Improved Loading vs Standard Loading | -0.2142 | 6.887 | Yes | ** | -0.3437 to -0.08465 |
| Improved Loading vs All Cell | -0.09417 | 3.028 | No | ns | -0.2237 to 0.03535 |
| Standard Loading vs All Cell | 0.12 | 3.61 | No | ns | -0.01846 to 0.2585 |

| Inv. AUC Percentage | | | | | |
|---|---|---|---|---|---|
| Tukey's Multiple Comparison Test | Mean Diff. | q | Significant? P < 0.05? | Summary | 95% CI of diff |
| Improved Loading vs Standard Loading | -0.2367 | 6.099 | Yes | ** | -0.3983 to -0.07506 |
| Improved Loading vs All Cell | -0.07667 | 1.976 | No | ns | -0.2383 to 0.08494 |
| Standard Loading vs All Cell | 0.16 | 3.857 | No | ns | -0.01277 to 0.3328 |

| Percent Counts Ambinet | | | | | |
|---|---|---|---|---|---|
| Tukey's Multiple Comparison Test | Mean Diff. | q | Significant? P < 0.05? | Summary | 95% CI of diff |
| Improved Loading vs Standard Loading | -12.46 | 5.13 | Yes | * | -22.57 to -2.344 |
| Improved Loading vs All Cell | -1.522 | 0.6266 | No | ns | -11.64 to 8.592 |
| Standard Loading vs All Cell | 10.94 | 4.213 | Yes | * | 0.1243 to 21.75 |

| Inv. Secant Line St.Dev. | | | | | |
|---|---|---|---|---|---|
| Tukey's Multiple Comparison Test | Mean Diff. | q | Significant? P < 0.05? | Summary | 95% CI of diff |
| Improved Loading vs Standard Loading | -0.06508 | 6.948 | Yes | ** | -0.1041 to -0.02607 |
| Improved Loading vs All Cell | -0.02642 | 2.82 | No | ns | -0.06543 to 0.01260 |
| Standard Loading vs All Cell | 0.03867 | 3.861 | No | ns | -0.003044 to 0.08038 |

| Inv. Cell's Scaled Slope Sum | | | | | |
|---|---|---|---|---|---|
| Tukey's Multiple Comparison Test | Mean Diff. | q | Significant? P < 0.05? | Summary | 95% CI of diff |
| Improved Loading vs Standard Loading | -0.3067 | 7.562 | Yes | ** | -0.4756 to -0.1378 |
| Improved Loading vs All Cell | -0.25 | 6.165 | Yes | ** | -0.4189 to -0.08110 |
| Standard Loading vs All Cell | 0.05667 | 1.307 | No | ns | -0.1239 to 0.2372 |

| Avg. Total Number of Cells | | | | | |
|---|---|---|---|---|---|
| Tukey's Multiple Comparison Test | Mean Diff. | q | Significant? P < 0.05? | Summary | 95% CI of diff |
| Improved Loading vs Standard Loading | 1861 | 4.98 | Yes | * | 304.4 to 3418 |
| Improved Loading vs All Cell | 2504 | 6.7 | Yes | ** | 947.4 to 4061 |
| Standard Loading vs All Cell | 643 | 1.609 | No | ns | -1021 to 2307 |

| Avg. Percent Counts Mitochondria | | | | | |
|---|---|---|---|---|---|
| Tukey's Multiple Comparison Test | Mean Diff. | q | Significant? P < 0.05? | Summary | 95% CI of diff |
| Improved Loading vs Standard Loading | 0.2208 | 0.2455 | No | ns | -3.525 to 3.967 |
| Improved Loading vs All Cell | 0.1908 | 0.2122 | No | ns | -3.555 to 3.937 |
| Standard Loading vs All Cell | -0.03 | 0.0312 | No | ns | -4.035 to 3.975 |

| Avg. Total Genes Detected | | | | | |
|---|---|---|---|---|---|
| Tukey's Multiple Comparison Test | Mean Diff. | q | Significant? P < 0.05? | Summary | 95% CI of diff |
| Improved Loading vs Standard Loading | -878.6 | 3.713 | No | ns | -1864 to 106.9 |
| Improved Loading vs All Cell | 328.2 | 1.387 | No | ns | -657.3 to 1314 |
| Standard Loading vs All Cell | 1207 | 4.771 | Yes | * | 153.3 to 2260 |

| Avg. Total Transcripts | | | | | |
|---|---|---|---|---|---|
| Tukey's Multiple Comparison Test | Mean Diff. | q | Significant? P < 0.05? | Summary | 95% CI of diff |
| Improved Loading vs Standard Loading | -4807 | 3.317 | No | ns | -10843 to 1229 |
| Improved Loading vs All Cell | -392.9 | 0.2711 | No | ns | -6429 to 5643 |
| Standard Loading vs All Cell | 4414 | 2.849 | No | ns | -2038 to 10867 |
