## Supplemental Table 1 for "A contamination focused approach for optimizing the single-cell RNA-seq experiment"

**Table S1 Quality Control Metrics on Experimental Data with Various Degress of Quality**

| | Contamination Metrics | | | | | Standard Metrics | | | |
|---|---|---|---|---|---|---|---|---|---|
| **Dataset** | **Empty Droplet's Scaled Slope Sum** | **Inverted Max. Secant Distance** | **Inverted Secant Line St. Dev.** | **Inverted AUC Percentage** | **Avg. Percent Counts Ambient** | **Total Number of Cells** | **Avg. Percent Counts Mitochondrial** | **Avg. Total Transcripts** | **Avg. Total Genes Detected** |
| **K562 Sample 1** | 0.17 | 0.28 | 0.282 | 0.20 | 5.50 | 239 | 9.62 | 28061.28 | 7330.51 |
| **K562 Sample 2** | 0.15 | 0.39 | 0.315 | 0.30 | 0.00 | 1248 | 0.65 | 1994.62 | 841.00 |
| **K562 Sample 3** | 0.11 | 0.34 | 0.301 | 0.26 | 0.00 | 1009 | 0.74 | 2362.86 | 897.48 |
| **Cold Protease on Crypts** | 0.61 | 0.69 | 0.405 | 0.60 | 18.26 | 1090 | 6.43 | 9395.47 | 3026.36 |
| **Gastric Corpus** | 0.75 | 0.86 | 0.459 | 0.80 | 56.89 | 1000* | 31.60 | 1542.61 | 453.61 |

*The total number of cells for the gastric corpus dataset was estimated as the ambient RNA level was too high to distinguish real cell and empty droplets
