## Supplemental Table 2 for "A contamination focused approach for optimizing the single-cell RNA-seq experiment"

**Supplementary Table 2 quality control metrics on total tissue dissociation under different loading conditions**

| | Contamination Metrics | | | | | Standard Metrics | | | |
|---|---|---|---|---|---|---|---|---|---|
| | Cell's Scaled Slope Sum | Max Secant Distance | Secant Line St.Dev. | AUC Percentage | Percent Counts Ambient | Avg. Total Number of Cells | Avg. Percent Counts Mitochondria | Avg. Total Genes Detected | Avg. Total Transcripts |
| Improved Loading on Total Tissue | 0.51 | 0.54 | 0.361 | 0.42 | 2.86 | 1032 | 4.11 | 2068.38 | 4809.52 |
| Wide Bore Loading on Total Tissue | 0.63 | 0.62 | 0.386 | 0.5 | 8.79 | 755 | 6.57 | 1843.19 | 4158.61 |
| MACS enzyme on Total Tissue with Standard Loading | 0.82 | 0.78 | 0.436 | 0.7 | 27.06 | 730 | 9.86 | 990 | 1860.7 |
| Cold protease on Total Tissue with Standard Loading | 0.78 | 0.83 | 0.450 | 0.76 | 22.34 | 295 | 5.06 | 1434.31 | 3220.45 |
